# Study of the glycome of synovial mast cells in normal tissues, osteoarthritis and rheumatoid arthritis reveals MGAT5 activity in these and endochondral cells

**DOI:** 10.1101/2022.05.19.492654

**Authors:** Sheena F McClure, John McClure

## Abstract

Mast cells and angiogenesis play roles in the synovium of both osteoarthritis and rheumatoid disease but precise mechanisms are unclear. In a study of the glycome of human synovial biopsies from normal, osteoarthritic and rheumatoid disease cases mast cell numbers were increased in both osteoarthritic and rheumatoid disease synovial samples (the latter more than the former). Both mast cells and endothelial cells expressed receptors for the lectin *Phaseolus vulgaris* leukagglutinin (lPHA). Endothelial cells only expressed receptors for *Psophocarpus tetragonolobus* (PTL-II). Mast cells and angiogenesis are, therefore, likely to have a common pathogenetic mechanism in osteoarthritis and rheumatoid synovitis, the difference being one of degree. lPHA receptor expression in both mast cells and endothelial cells indicates βl,6N–acetylglucosaminyl (GlcNAc) – transferase V (GnTaseV/MGAT5) activity. This provides a linkage between mast cells and endothelial cell proliferation. Endothelial cells show core 1 O-glycosylation (positive PTL-II staining) which is probably necessary for the formation of competent tubular structures and for continuing vascular integrity.

## INTRODUCTION

The glycome is the complete set of glycans and glycoconjugates made by a cell or organism (Hart & Copeland, 2010). These macromolecules have a high information content and their formation is probably the most important post-translational event in determining phenotypic variation. The importance of the glycome is increasingly recognised (Reily et al. 2019). Although exploration of the glycome has been technically difficult, the use of lectin histochemistry can provide insights (Barkhordari et al. 2004).

In addition to its role in allergic reactions, the ubiquitous mast cell has been implicated in angiogenesis, lymphangiogenesis, tumour growth and metastasis (Aponte-Lopez at al. 2018). However mast cells exhibit considerable phenotypic heterogeneity and may have both agonistic and antagonistic effects (Varrichi et al. 2017).

The fact of increased numbers of mast cells in the synovial tissues in rheumatoid arthritis (RA) is well established (Athreya et al. 1978; Crisp 1984; Crisp et al. 1984; Godfrey et al. 1984; Gruber et al. 1986). Conventionally, osteoarthritis (OA) has been attributed to cartilage wear. However, there is emerging evidence of an inflammatory process and it is likely that mast cells play a key role in this. Mast cell numbers in OA have been reported as greater than in synovial controls (Buckley et al, 1998) and higher than in rheumatoid synovia (deLange-Brokar, 2016). Gigante et al (2016) reviewed the literature concluding that mast cells play a complicated multifaceted role in the pathogenesis of OA. Wang et al. (2019) have produced evidence that IgE-mediated mast cell activation promotes inflammation and cartilage destruction in OA.

Despite evidence of its involvement in a number important diseases little is known about the mast cell glycome. The present study was a detailed lectin histochemical study of the mast cell glycome designed to elucidate its role in normal, osteoarthritic and rheumatoid synovia.

## METHODS

Paraffin-embedded tissue blocks of synovium from normal, osteoarthritic and rheumatoid arthritic joints taken by biopsy were obtained from a tissue archive. Cases were selected on the basis of there being sufficient material in the blocks to allow a full range of the histological, immunohistological and lectin histochemical studies to be performed. Independently, the cases were given a randomly generated anonymising identifier number so that all tests and assessments were made without knowledge of patient clinical and demographic details. After decodification and assignment of cases to their particular categories, comparative analyses were performed.

In the group of normals there were nine females and seven males with a mean age of 40.5 years (range 14 – 69). Samples were from 13 knee, one hip, one finger and one elbow joints. The osteoarthritic (OA) samples were from nine knee and three hip joints. There were nine males and three females with a mean age of 64.6 years (range 39 – 84). For rheumatoid arthritic (RA) patients the mean age was 60.0 years (range 38 – 78) for 10 females and two males. Samples were from five knee, three wrist, two finger, one hip and one elbow joints.

Sections (5 μm) were cut from all blocks and stained with haematoxylin and eosin (H & E) for histological examination and by a conventional immunological technique for mast cell tryptase to detect connective tissue mast cells. In addition, sections were stained with a panel of lectins. These are listed in Table 1 showing their group, origin and carbohydrate specificity.

**TABLE 1.**
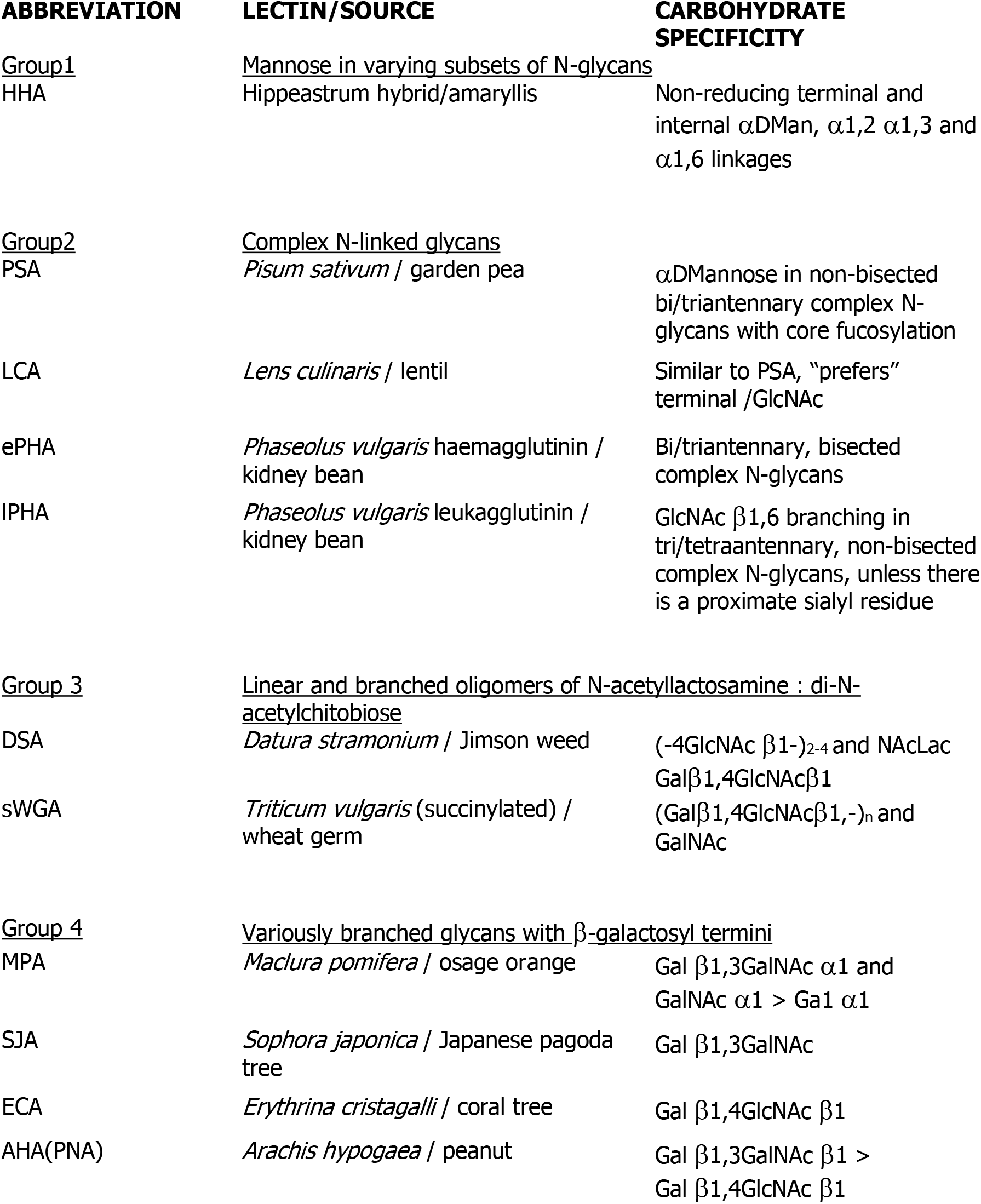
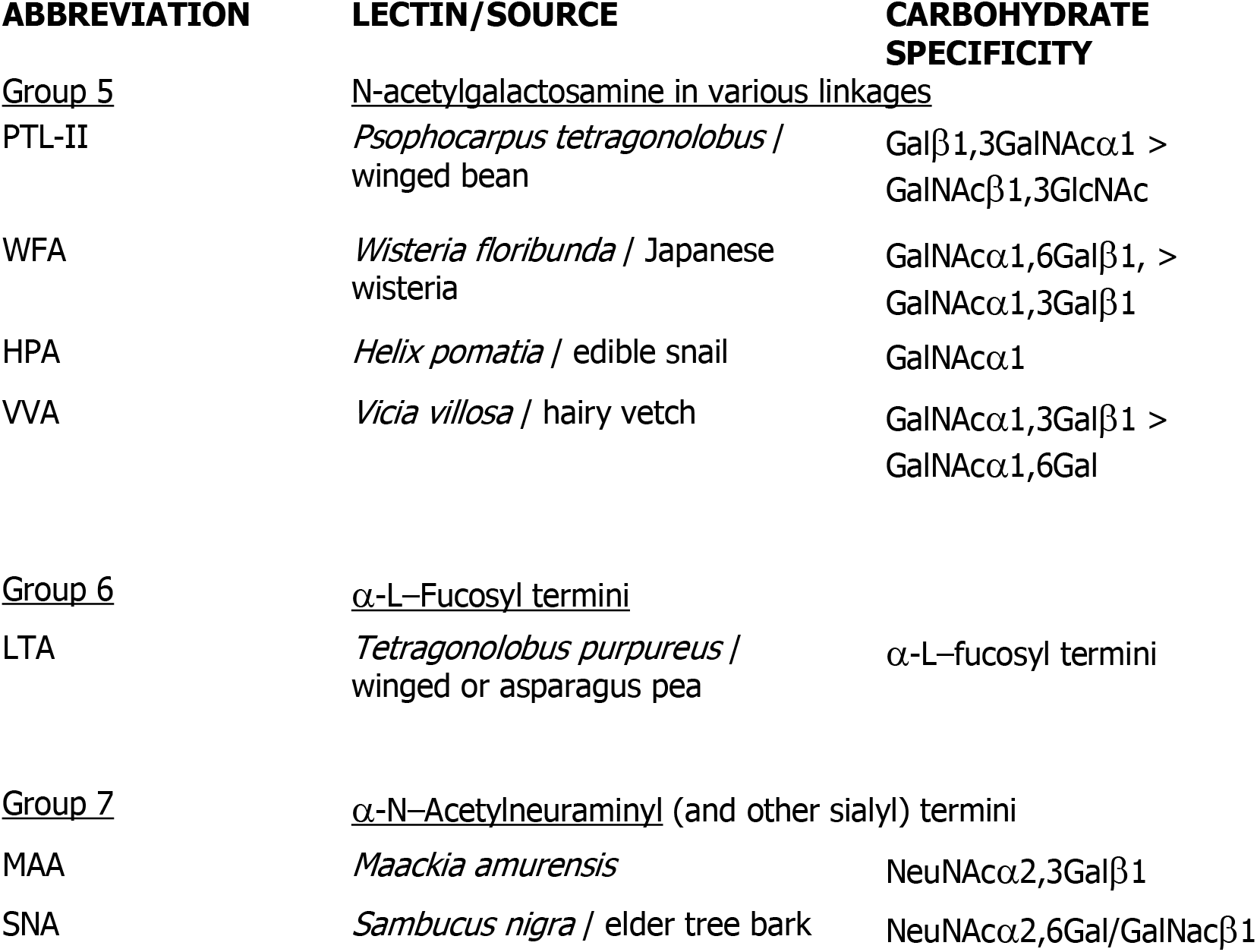
The group origin and carbohydrate specificity of the lectins used in the study are shown.

Sections were stained with biotinylated lectins following treatment with crude trypsin (Jones and Stoddart 1986). Sections were treated with lectins at a concentration of 10 μg/ml followed by avidin-conjugated peroxidase (5 μg/ml in 0.125 M TBS, pH 7.6, containing 0.374 M sodium chloride), then 3,3–diaminobenzidine tetrahydrochloride and counterstained with methyl green. The controls for the lectin histochemistry included negative controls with buffer substitution for the lectin, competing sugars and β-elimination for MAA and SNA.

All sections were examined for the presence of positive lectin staining reactions. Mast cell counts were performed on sections stained for mast cell tryptase. Blood vessels stained by PTL-II were also counted (a complete lumen was required for inclusion) and in both instances counts were performed on the screen of a Nikon Coolscope at a magnification of 200x and 10 fields were recorded for each case.

## RESULTS

Of the 18 lectins applied synovial mast cells stained with 13 and five of these showed cell membrane staining only (PSA/LCA/ECA/SWGA/MAA-II). The remaining eight showed generalized cytoplasmic staining. Nuclear staining was never observed. Table 2 shows the percentage of cases in which staining of mast cells by a particular lectin was observed. As shown, there were a number of cases in which staining was never observed despite assiduous searching. Universally across the three groups (normals, OA, RA) mast cells were stained by HHA. Seven lectins (LCA/ePHA/DSA/MPA/SWGA/MAAII/SNA) showed some but relatively few changes in staining between the three groups. Five lectins (PSA/lPHA/ECA/WFA) showed greater changes. For PSA, OA and RA showed increased percentages of cases staining compared to normals. The same changes were shown for lPHA. For ECA, OA percentage was reduced compared to normals and RA. With WFA, OA was reduced compared to normals and RA became zero. For AHA, RA was reduced compared to normals and OA.

**TABLE 2.** Percentage of cases in each group (N, OA, RA) in which mast cells showed positive staining reactions with individual lectins.

|  | **N** | **OA** | **RA** |
| --- | --- | --- | --- |
| **Group 1** | **Mannose in varying subsets of N-glycans** |  |  |
| HHA | 100% | 100% | 100% |
| **Group 2** | **Complex N-linked glycans** |  |  |
| PSA | 44% | 75% | 75% |
| LCA | 94% | 83% | 92% |
| ePHA | 88% | 92% | 83% |
| IPHA | 44% | 92% | 100% |
| **Group 3** | **Linear and branched oligomers of N-acetyl lactosamine : di-N-acetylchitobiose** |  |  |
| DSA | 81% | 92% | 75% |
| sWGA | 69% | 83% | 67% |
| **Group 4** | **Variously branched glycans with β-galactosyl termini** |  |  |
| MPA | 75% | 83% | 75% |
| SJA | 0 | 0 | 0 |
| ECA | 69% | 42% | 58% |
| AHA | 75% | 75% | 25% |
| **Group 5** | **N-acetylgalactosamine in various linkages** |  |  |
| PTL-II | 0 | 0 | 0 |
| WFA | 50% | 17% | 0 |
| HPA | 0 | 0 | 0 |
| VVA | 0 | 0 | 0 |
| **Group 6** | **α-L-Fucosyl termini** |  |  |
| LTA | 0 | 0 | 0 |
| **Group 7** | **α-N-Acetylneuraminyl (and other sialyl) termini** |  |  |
| MAA-II | 81% | 92% | 83% |
| SNA | 81% | 92% | 75% |

Table 3 shows mast cell and blood vessel counts for normals, OA and RA cases.

**TABLE 3.** Mast cell and blood vessel counts in normal, OA and RA synovial samples.

|  | **Mean** | **Standard Deviation** |
| --- | --- | --- |
| Normals (N) | 1.500 | 0.861 |
| OA cases (OA) | 6.233 | 3.014 |
| RA cases (RA) | 17.250 | 4.738 |

**BLOOD VESSEL COUNTS**
|  | **Mean** | **Standard Deviation** |
| --- | --- | --- |
| Normals (N) | 7.667 | 2.440 |
| OA cases (OA) | 16.500 | 7.380 |
| RA cases (RA) | 52.450 | 23.793 |

N<OA (p<0.0001); N<RA (p<0.0001); OA<RA (p<0.0001) (Wilcoxon signed rank test)

The lectin PTL-II stained only the cytoplasm of capillary blood vessels of all cases in the three groups. Identical, but less intense staining, was obtained with lPHA.

## DISCUSSION

The results of the present study confirm that mast cell numbers are significantly higher in rheumatoid synovium compared to both normals and osteoarthritic synovium. The latter contains mast cell numbers also significantly greater than normals. It is known that mast cell numbers are increased in osteoarthritic synovium and synovial fluid and it has been speculated that there are common pathogenetic mechanisms with rheumatoid synovitis (Dean et al. 1983).

The precise role of mast cells in such mechanisms remains undetermined. These cells contain a large number of mediators and possible actions include recruitment of inflammatory cells, induction of synoviocyte hyperplasia and further mediator production, stimulating the production of degradative enzymes and fostering angiogenesis (Nigrovic and Lee 2004).

There are relatively few published papers describing lectin binding studies of human mast cells. Data are particularly sparse in relationship to truly normal tissues and extended lectin panels have not been applied often. Generally mannose and complex N-glycans have been detected and, indeed, this is the case in the present study (Schumacher et al. 1987; Kirkpatrick et al.1988; Schumacher et al. 1991). In contrast O-glycans have not been reported whereas in the present study staining reactions to lectins in groups 3, 4 and 5 indicate their presence. As in the present study, fucosylation has never been reported. Positive reactions with MAA, and SWA indicate the presence of sialylated structures. In an analysis of human skin mast cell proteins by two-dimensional gel electrophoresis tryptase was identified as a sialylated glycoprotein (Benyon et al. 1993). Sialic acid has been identified in a human skin mastocytoma (Schumacher et al. 1989).

It is clear from the present study that whilst the mast cell glycome is restricted there are some variations between cells in normal, OA and RA synovia. The significance of these variations in terms of changes in the chemical structures detected can be assessed by reference to the relevant carbohydrate specificities. However, it is usually not possible to assess these variations in terms of functional significance. Nevertheless, the observations underscore the increasingly recognized heterogeneity of mast cells. Final maturation of the cells occurs after the arrival of blood borne progenitor cells in the target tissue. Local heterogeneity in terms of size, granule content and mediator content is in response to local signals (Metcalf et al. 1997). The results of the present study clearly demonstrate heterogeneity of glycans expression in relationship to disease.

A lectin from group 5 (PTL-II) did not stain mast cells but specifically and universally stained endothelial cells only. This staining was intense, diffuse and cytoplasmic in location. Endothelial cells also stained with lPHA. Again the staining was cytoplasmic but more coarsely granular. The stained endothelial cells were in the synovial subserosa. Counts of these vessels showed significantly greater numbers in RA synovia compared to normals and OA and significantly greater numbers in RA versus OA synovia. Angiogenesis is occurring in both OA and RA synovia and is significantly greater in the latter.

Group 5 lectins combine with N-acetylgalactosamine in various linkages. Specifically the receptor for PTL-II is Galβ1,3GalNAcα1>GalNAcβ1,3GlcNAc. The responsible enzyme for the formation of this core 1 O-glycans is Galβ1,3GalNacα1 is β1,3 galactosyltransferase, also known as T synthase. Core 1 O-glycans in wild type mice is expressed primarily in endothelial, haemopoietic and epithelial cells during development (Xia et al. 2004). Gene-targeted mice lacking T synthase suffered brain haemorrhage which was inevitably fatal by embryonic day 14. T synthase deficient mice brains had a chaotic microvascular network with distorted capillary lumina and defective association of endothelial cells with pericytes and extracellular matrix. These observations revealed an unexpected requirement for core 1 O-glycans during angiogenesis and that requirement had to do with proper structural organisation of the blood vessel at the microscopical level. It has been suggested that altogether there is a conserved role in the molecular events governing tubulogenesis across diverse organs and species possibly by influencing trafficking and stability of crucial proteins involved in tube architecture and function (Tian and Ten Hagen, 2009). It would appear that O-glycosylation is a feature of human synovial angiogenesis and is presumably necessary for the development and maintenance of vascular tubular structures. There is also evidence that O-glycosylation maintains the separation of blood and lymphatic vessels throughout life and preserves vascular integrity (Herzog et al. 2014).

There is a body of opinion that angiogenesis is important in the initiation and progression of rheumatoid arthritis (Koch 1998; Paleolog 2002). It has been pointed out that in the RA synovium there is active endothelial cell proliferation consequent to the synovium being an hypoxic environment. Expansion in the volume of the synovial membrane leads to a compensatory increase in the number and density of blood vessels. Fibroblast growth factor 2 (FGF-2) which has been described as the classical angiogenesis factor is a known constituent of synovial fluid in RA and its concentration is positively related to severity of clinical disease (Vlodavsky et al. 1987; Manabe et al. 1999).

There is substantial evidence linking mast cells and angiogenesis. Mast cells accumulate in a number of angiogenesis dependant processes such as wound healing and tumour growth. Several mast cell mediators (TNF, VEGF, FGF-2) are angiogenic. In addition, mast cell products such as tryptase can degrade connective tissue matrix providing space for neovascular sprouts (Hiromatsu and Toda 2003).

In the present study binding sites for lPHA were expressed by both synovial mast cells and endothelial cells. The percentage of cases in which there was expression of these binding sites by mast cells rose substantially from normal to OA and RA tissues. Positive reactions for endothelial cells were less extensive in normal samples. Generally group 2 lectins bind to complex N-glycans and where these form branching structures (antennae) lPHA specifically detects increased β1,6GlcNAc branching. This increased GlcNAcβ1,6Manα1,6Manβbranching in asparagine-linked glycans has been associated with enhanced metastatic potential of carcinomas and neo-angiogenesis is considered a likely factor (Dennis et al. 1987). Further it has been shown that lPHA requires this β1,6 branching for high-affinity binding and that β1,6 branching in asparagine-linked glycans is dependent on β1,6N-acetylglucosaminyl (GlcNAc) – transferase V (also known GnTase V or MGAT5) (Dennis and Laferte 1989). A modern view is that MGAT5 selectively remodels the endothelial cell surface by the regulated binding of Galectin-1 (Gal-1) which on recognition of complex N-glycans on VEGFR2 activates VEGF signalling (Croci et al. 2014). In experimental animals with complete absence of MGAT5, N-glycans on VEGFR2 bind insufficient Gal-1 to promote angiogenesis and thus tumour growth decreases (Stanley, 2014).

The expression of MGAT5 by both mast cells and endothelial cells in synovial tissues is conceivably related to angiogenesis in the arthritides. The first by an exocrine action and the second by an autocrine one.

---

**Figure 1.**
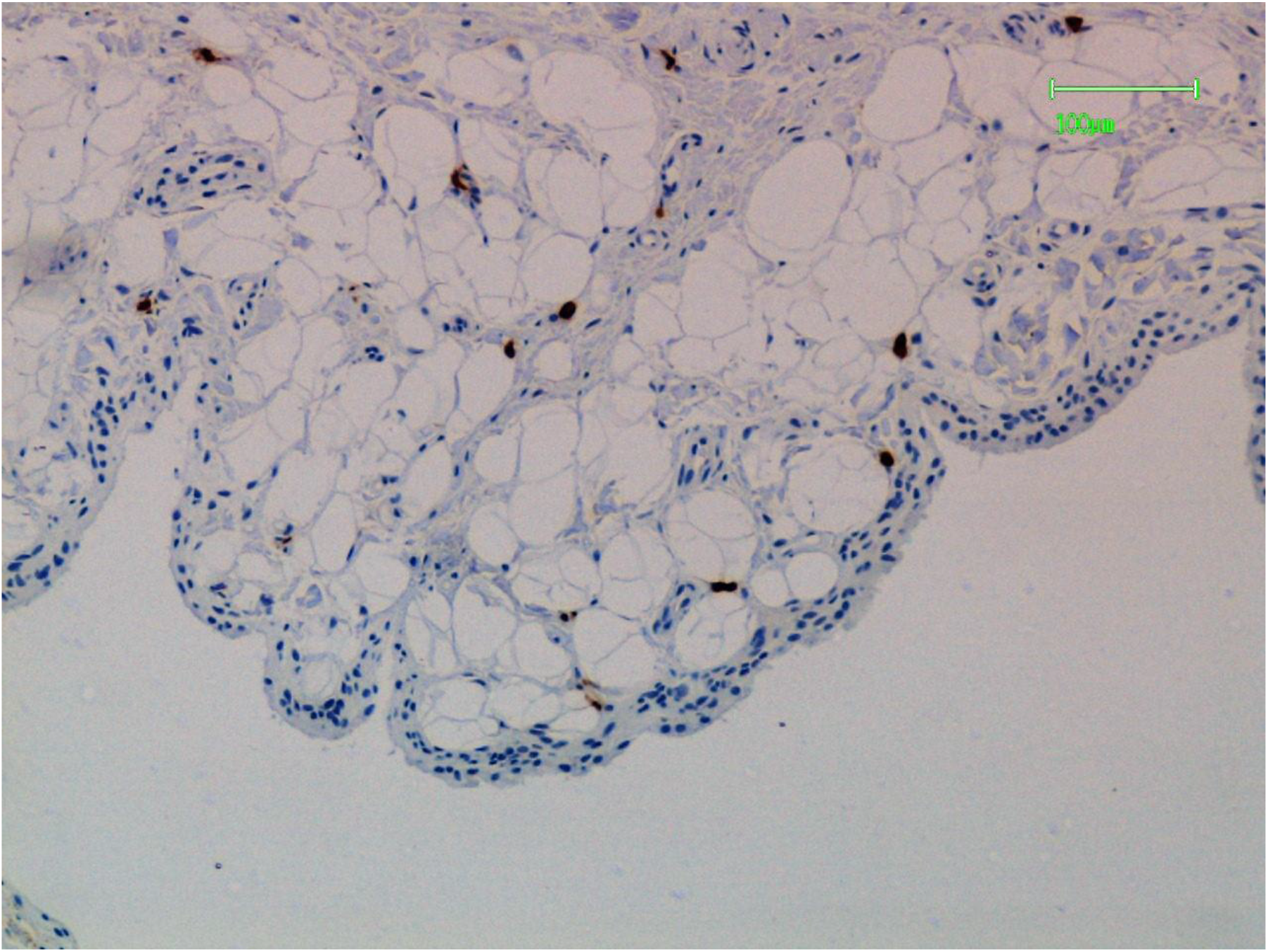
This is a sample of normal synovium. Mast cells have been stained for tryptase (brown deposit). They are located below the surface layer and throughout the fibrofatty stroma.

**Figure 2.**
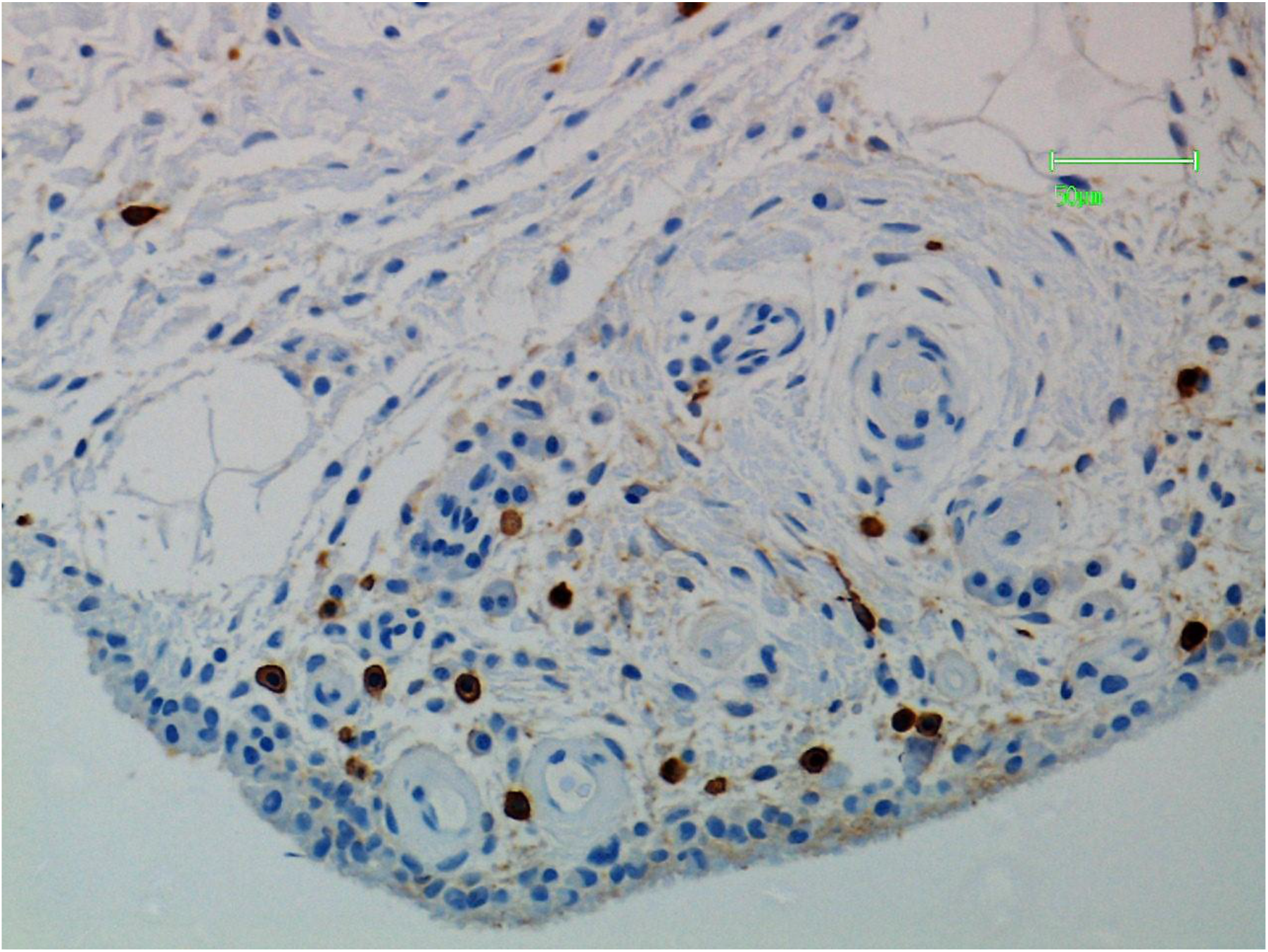
This is a sample of synovium in osteoarthritis (OA). Mast cells have been stained for tryptase (brown deposit). They are located in the stroma below the surface layer.

**Figure 3.**
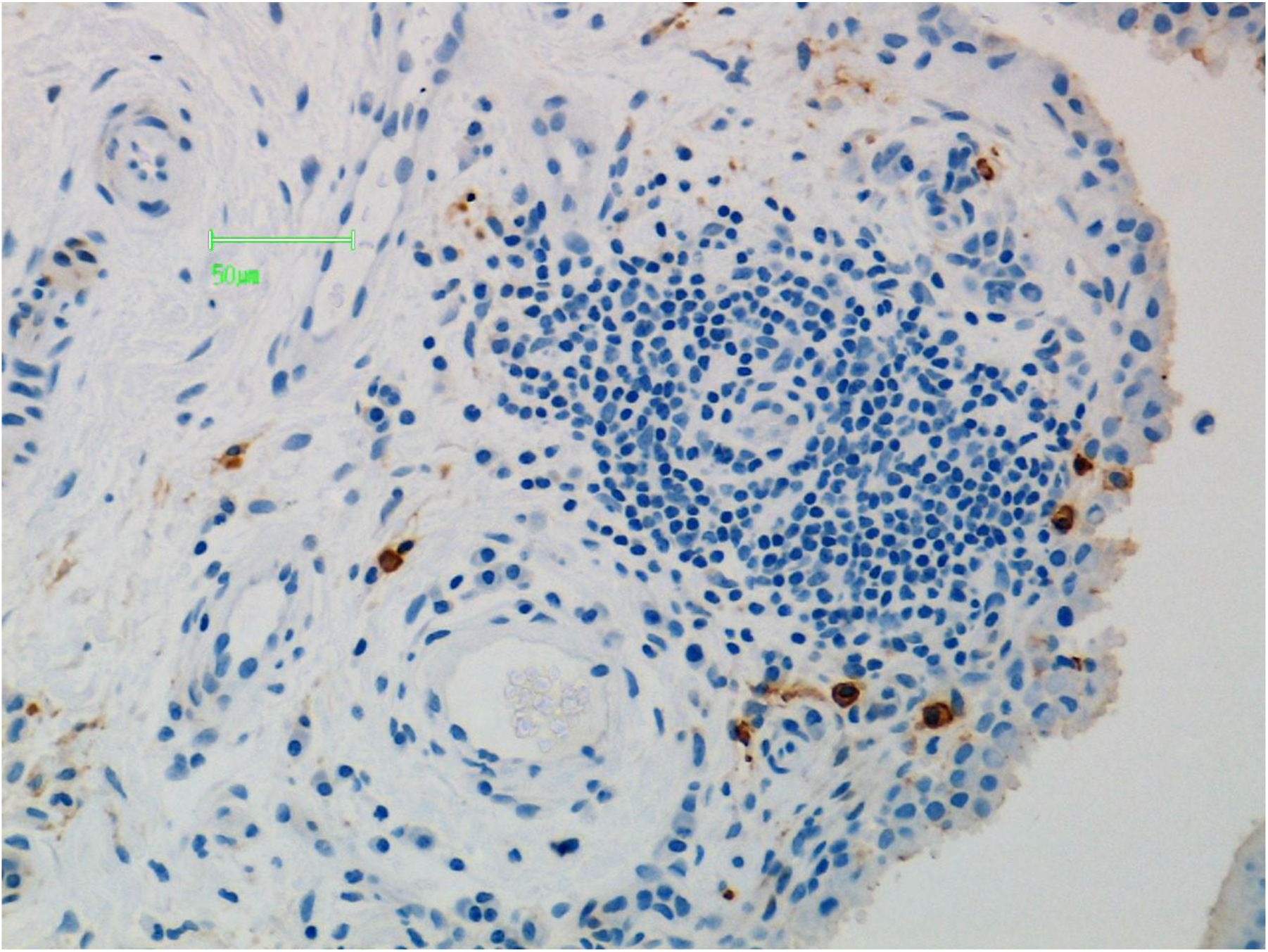
This is a sample of synovium in rheumatoid arthritis (RA). Mast cells have been stained for tryptase (brown deposit). Some are infiltrating the surface layer. Others in the stroma are adjacent to blood vessels.

**Figure 4.**
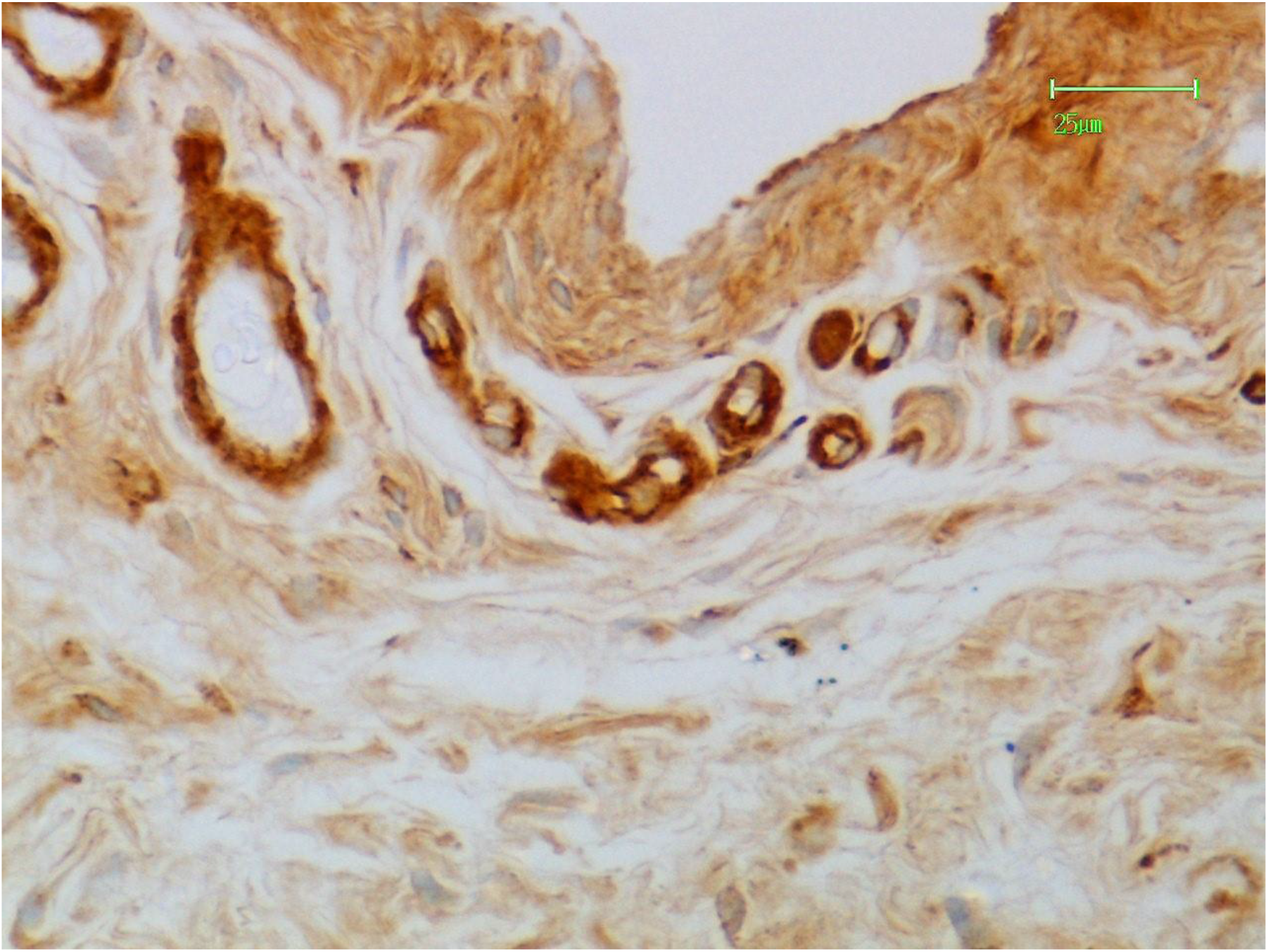
This is a sample of synovium in osteoarthritis (OA). The tissue has been stained by lPHA lectin. Positive reactions (brown deposit) are present the surface layer in cell cytoplasm. Stronger reactions are present in mast cells and blood vessel walls.

**Figure 5.**
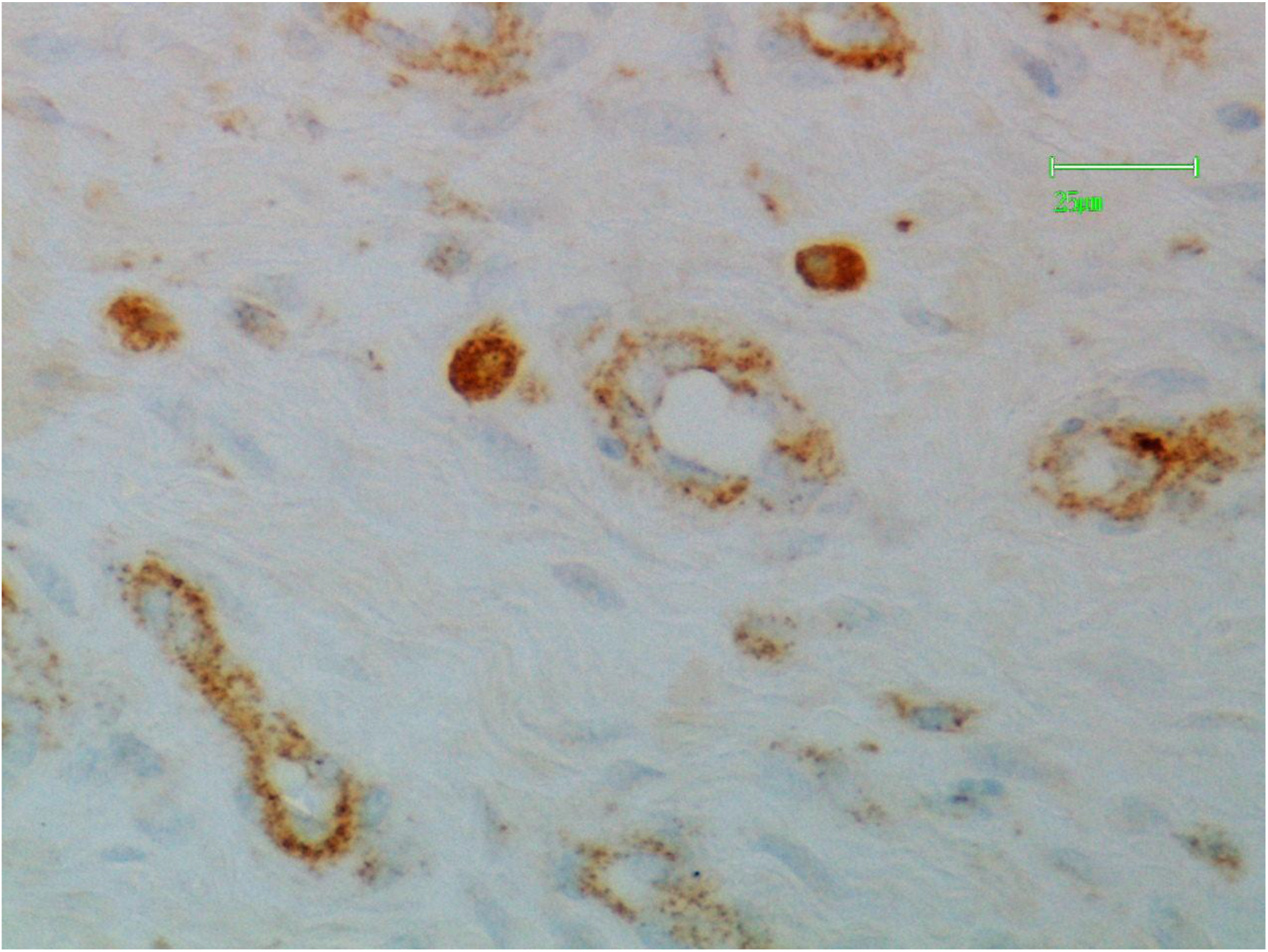
This is a sample of synovium in rheumatoid arthritis (RA). The tissue has been stained by lPHA lectin. There are granular brown positive reactions in the cytoplasm of mast cells and endothelial cells of the stroma.

**Figure 6.**
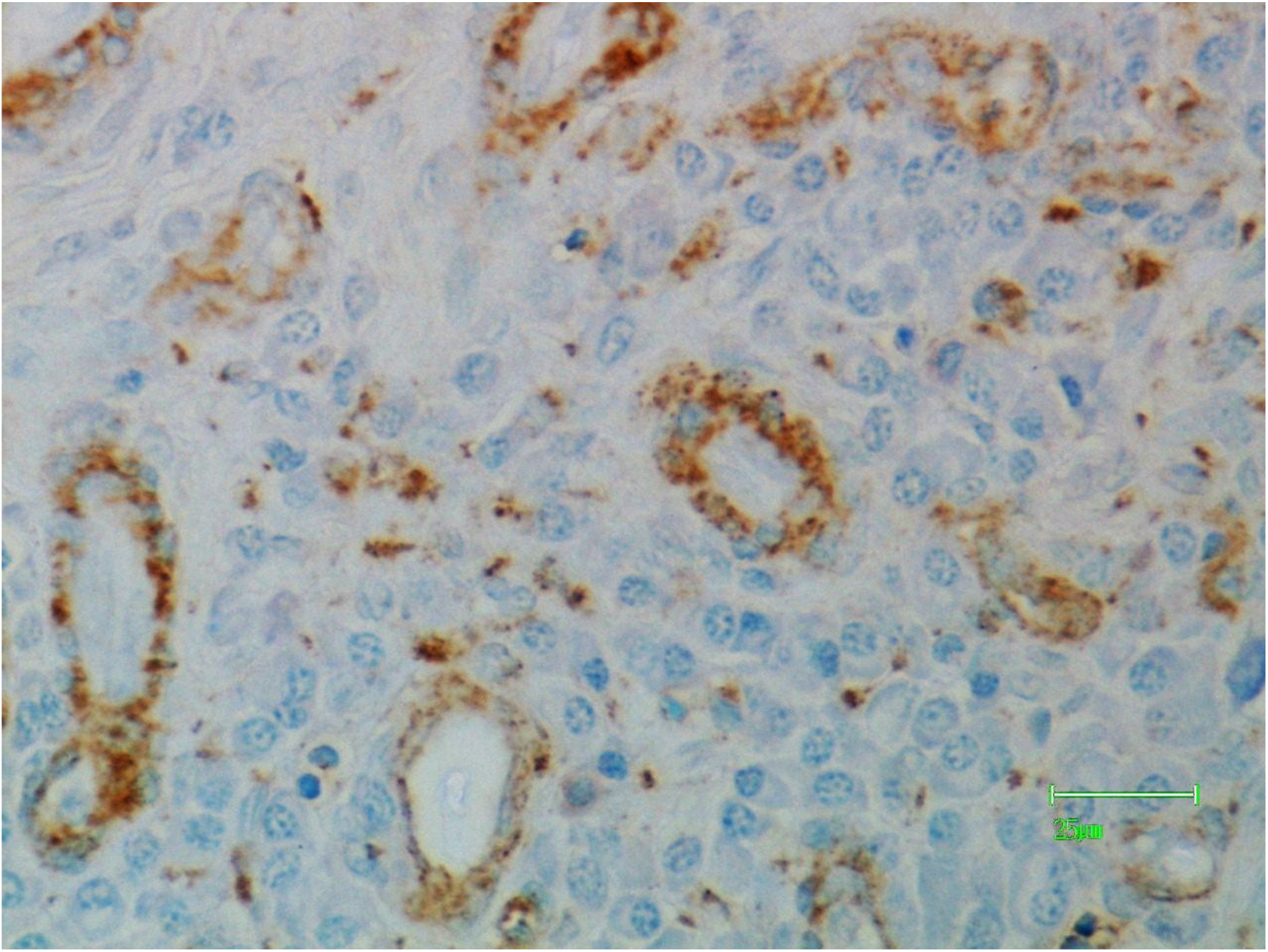
This is a sample of synovium in rheumatoid arthritis (RA). The tissue has been stained by lPHA lectin. Granular brown positive reactions are present in the cytoplasm of endothelial cells.

**Figure 7.**
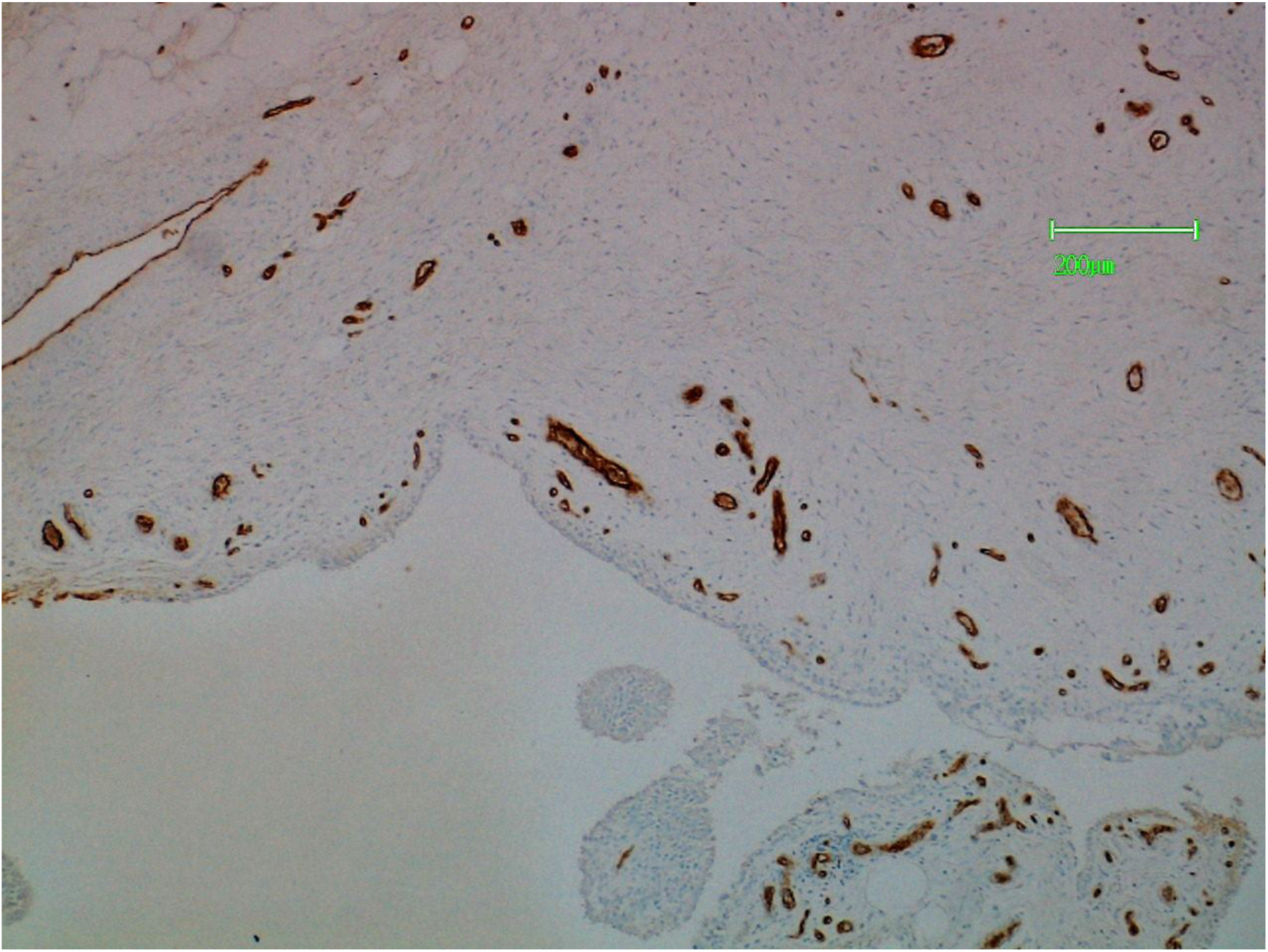
This is a sample of normal synovium stained by PTL-II lectin. Blood vessels throughout the stroma show positive reactions.

**Figure 8.**
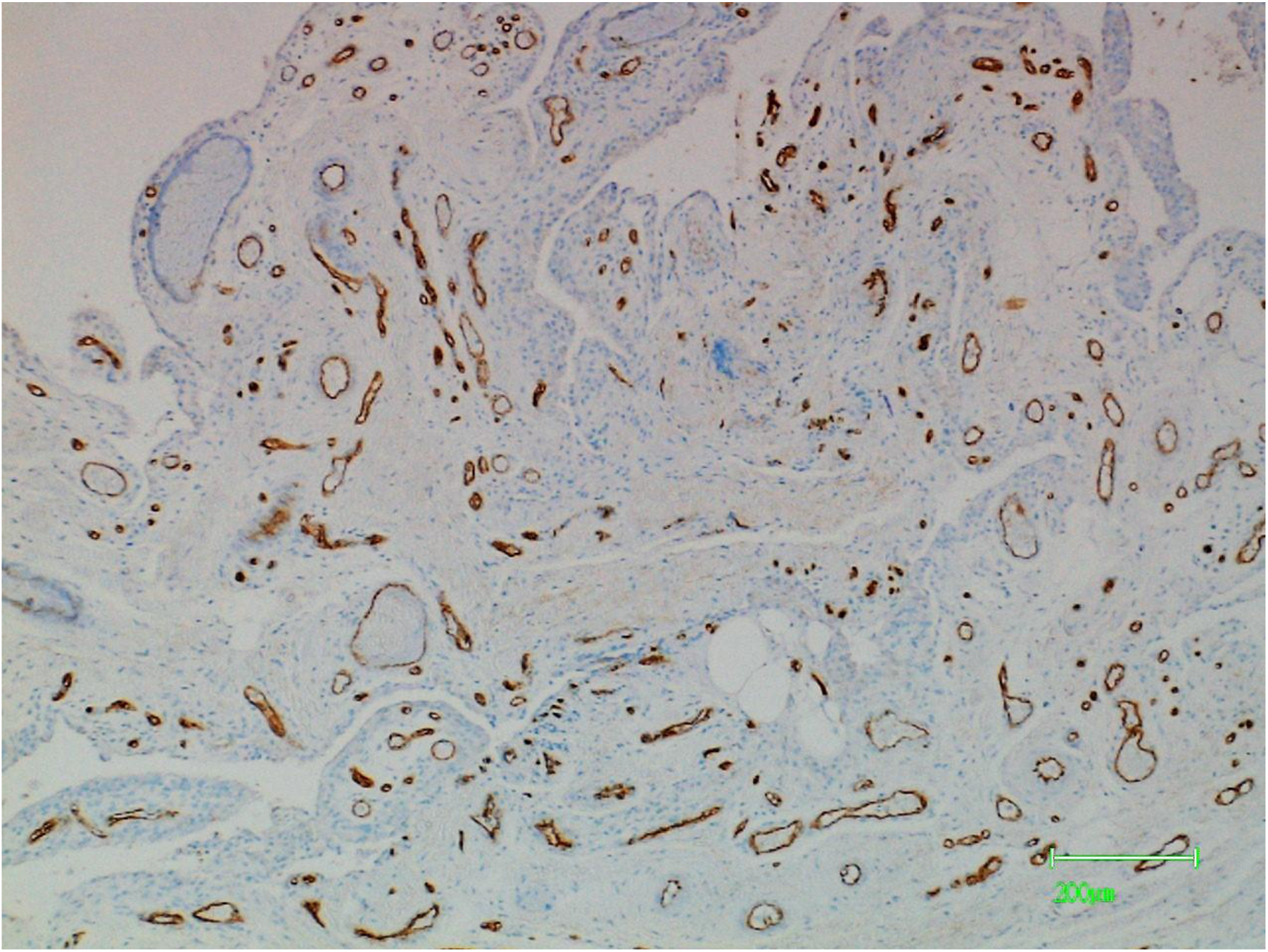
This is a sample of synovium in osteoarthritis (OA). It has been stained by PTL-II lectin with positively reacting blood vessels throughout the stroma.

**Figure 9.**
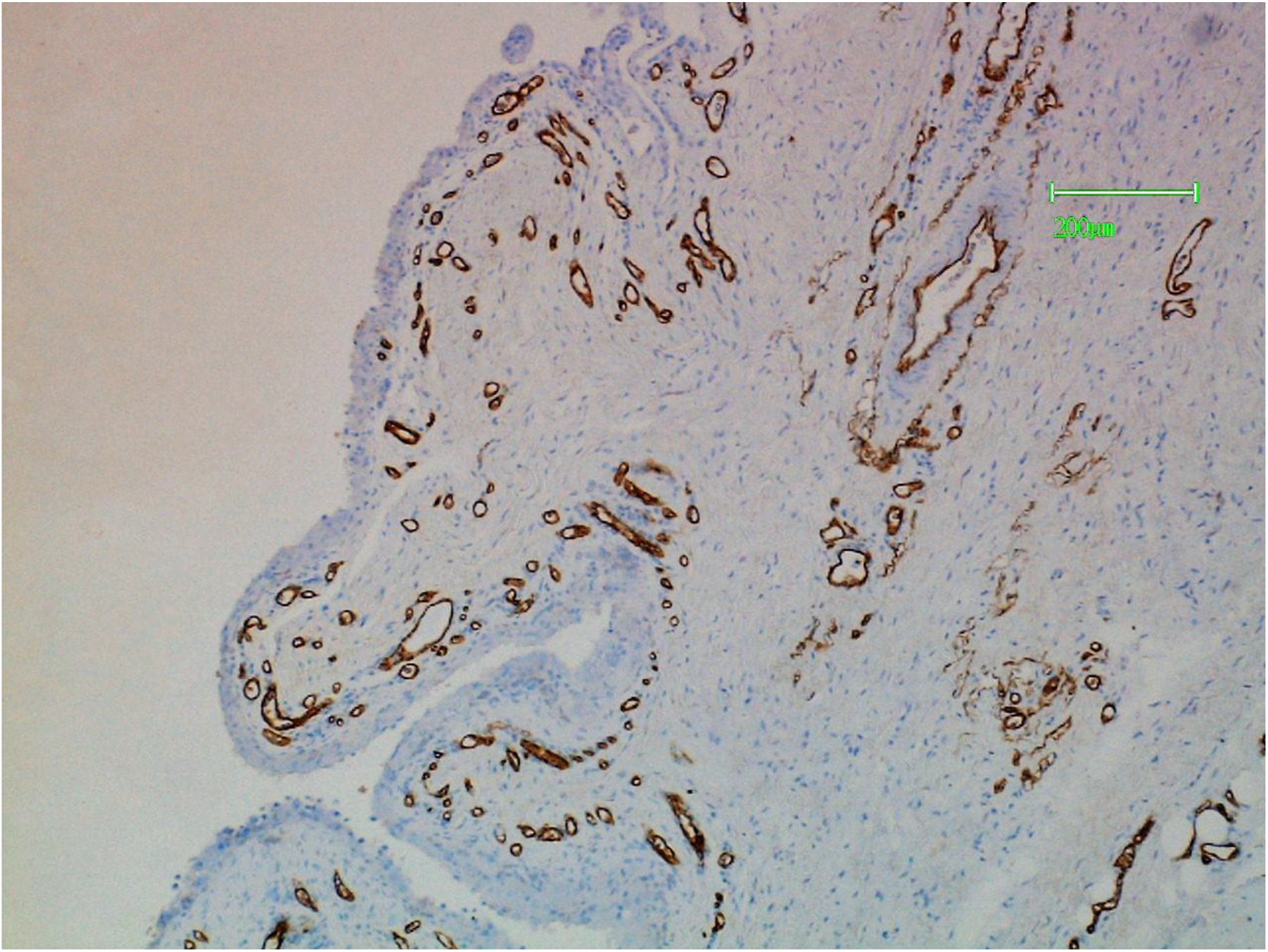
This is a sample of synovium in rheumatoid arthritis (RA). It has been stained by PTL-II lectin showing positively reacting blood vessels. In this section these are mainly located below the surface layer which shows hyperplasia.

**Figure 10.**
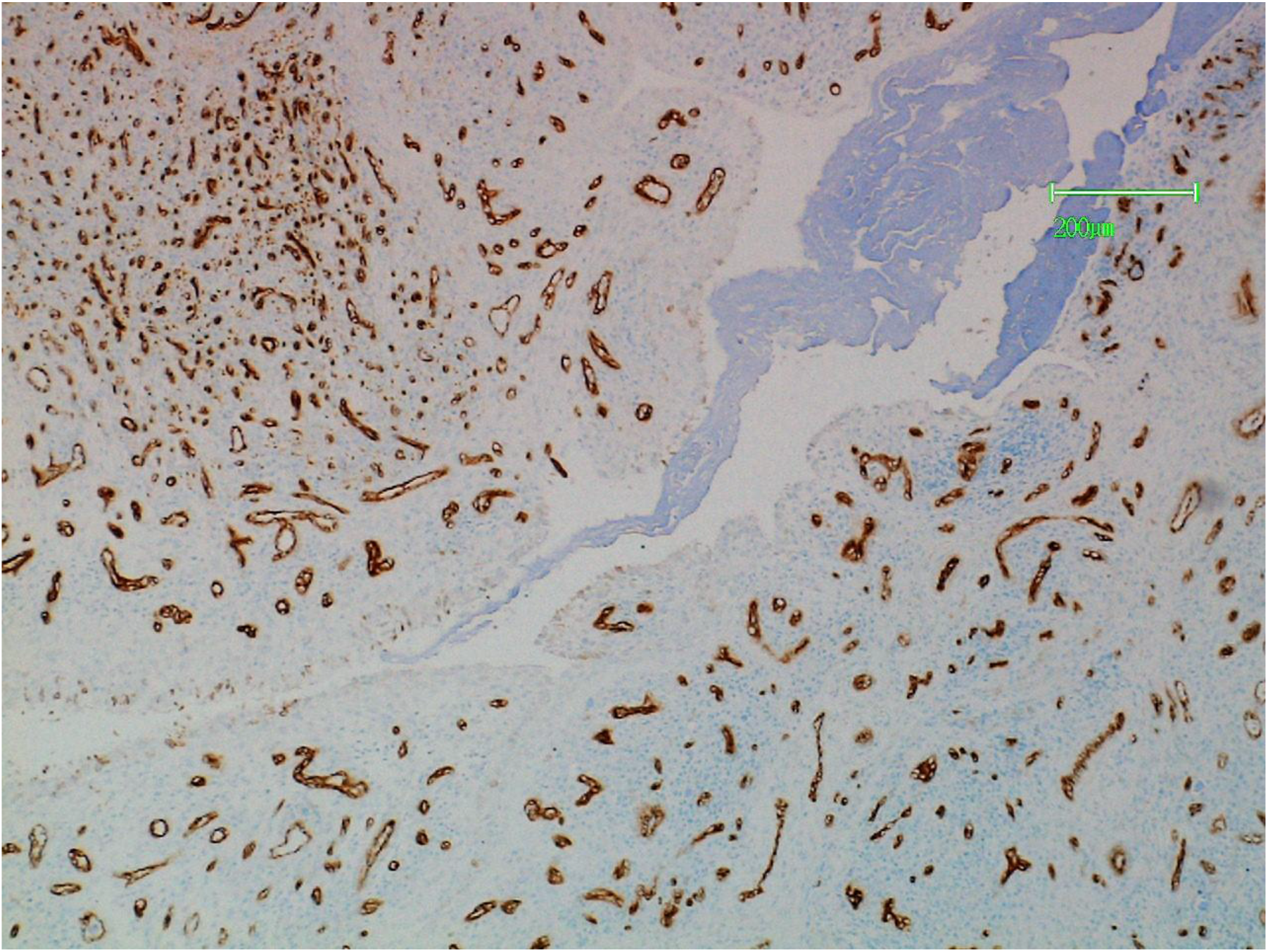
This is a sample of synovium in rheumatoid arthritis (RA). It has been stained with PTL-II lectin. The positively reacting blood vessels are distributed mainly in the stroma.

## REFERENCES

Aponte-Lopez A, Fuentes-Panana E, Cortez-Munoz D, Munoz-Cruz S. (2018) Mast cell, the neglected member of the tumour microenvironment. J Immunol Res: Article ID 2584243.

Athreya BH, Moser G, Schumacher HR, Hanson V, Dahms B, Thompson W. (1978) Role of basophils and mast cells in juvenile rheumatoid arthritis. In Pepys J. Edwards Am(eds) The Mast Cell: Its Role in Health and Disease. Pitman, New York, pp 127–136

Barkordari A, Stoddart RW, McClure SF, McClure J. (2004) Lectin histochemistry of normal human lung. J Mol Histol 35:147–56

Benyon RC, Enciso JA, Befus AD (1993) Analysis of human skin mast cell proteins by two-dimentional gel electrophoresis. Identification of tryptase as a sialylated glycoprotein. J Immunol 151:2699–2706

Buckley MG, Gallagher PJ, Walls AF. (1998) Mast cell populations in the synovial tissue of patients with osteoarthritis: selective increase in numbers of tryptase-positive, chymase-negative mast cells. J Pathol 186: 67–74

Crisp AJ (1984) Mast cells in rheumatoid arthritis. J Roy Soc Med 77:450–1

Crisp AJ, Chapman CM, Kirkham SE, Schiller AL, Krane SM (1984) Articular mastocytosis in rheumatoid arthritis. Arthritis Rheum 27:845–851

Croci DO, Cerliani JP, Dolatto-Moreno T, Mendez-Huergo SP, Mascanfroni ID et al (2014) Glycosylation-dependent lectin-receptor interactions preserve angiogenesis in anti-VEGF refractory tumors. Cell 156:744–758.

Dean G, Hoyland JA, Denton J, Donn RP, Freemont AJ (1993) Mast cells in the synovium and synovial fluid in osteoarthritis. Rheumatology 32:271–275

De Lange-Broker BJE et al. (2016) Characterisation of synovial mast cells in knee osteoarthritis: association with clinical parameters. Osteoarthritis Cartilage 24: 664–71.

Dennis JW, Laferte S, Waghorne C, Britman ML, Kerbel RS (1987) Beta 1-6 branching of Asn-oligosaccharide is directly associated with metastasis. Science 236:582–585

Dennis JW, Laferte S (1989) Oncodevelopmental expression of-GIcNAcβl-6Manα1-6Manβ1-branched asparagine-linked oligosaccharides in murine tissues and human breast carcinomas. Cancer Res 49:945–950

Gigante A, Farinelli L, Manzotti S, Aquili A, Papalia G (2018) Synovial mast cells in osteoarthritis. Osteoarthritis: Med Docs e Books 1–7.

Godfrey HP, Llardi C, Engber W, Graziano FM (1984) Quantitation of human synovial mast cells in rheumatoid and other rheumatic diseases. Arthritis Rheum 27:852–856

Gruber B, Poznansky M, Boss E, Partin J, Gorevic P, Kaplan AO (1986) Characterization and functional studies of rheumatoid synovial mast cells: activation by secretogogues, anti IgE and a histamine releasing lymphokine. Arthritis Rheum 29:944–955

Hart GW, Copeland RJ (2010) Glycomics hits the big time. Cell 143:672–676

Herzog BH et al. (2014) Mucin-type O-glycosylation is critical for vascular integrity. Glycobiology 27: 1237–41.

Hiromatsu Y, Toda S (2003) Mast cells and angiogenesis. Microsc Res Tech 60:64–69

Jones CJ, Stoddard RW (1986) A post-embedding avidin-biotin peroxidase system to demonstrate the light and electron microscopic localisation of lectin binding sites in rat kidney tubules. Histochem J 18:371–379

Kirkpatrick CJ, Jones CJ, Stoddard RW (1988) Lectin histochemistry of the mast cell: a light microscopical study. Histochem J 20:139–146

Koch AE (1998) Angiogenesis - Implications for rheumatoid arthritis. Arthritis Rheum 41:951–962

Manabe N, Oda H, Nakamura K, Kuga Y, Uchida S and Kawaguchi H (1999) Involvement of fibroblast growth factor in joint destruction of rheumatoid arthritis patients. Rheumatology 38:714–720

Metcalfe DD, Baram D, Mekor YA (1997) Mast cells. Physiol Rev 1997, 77:103–179

Nigrovic PA, Lee DM (2004) Mast cells in inflammatory arthritis. Arthritis Res Ther 7:1–11

Paleolog EM (2002) Angiogenesis in rheumatoid arthritis. Arthritis Res 4:581–590

Reily C, Stewart TJ, Renfrow MB, Novak J (2019) Glycosylation in health and disease. Nature Reviews 15:347–366

Schumacher U, Horny HP, Welsch U (1987) The lectin leucoagglutinin binds specifically to human granulocytes, monocytes and tissue mast cells: further evidence of a common origin of the three cell types. Br J Haematol 66:405–406

Schumacher U, Horny HP, Welsch U, Kaiserling E (1989) Lectin binding studies of a human skin mastocytoma. J Histochem 21:44–46

Schumacher U, Horny HP, Welsch U, Geerts ML, Kaiserling E (1991) Lectin binding patterns of human tissue mast cells indicated marked phenotypical diversity. Acta Histochem 91:141–146

Stanley P (2014) Galectin-1 pulls the strings on VEGFR2. Cell 156:625–626

Tian E, Ten Hagen KG (2009) Recent insights into the biological roles of mucin type O-glycosylation. Glycoconj J 26:325–334

Varrichi G, Galdiero MR, Loffedo S et al (2017) Are mast cells MASTers in cancer? Front Immunol 8:1–13

Vlodavsky I, Folkman J, Sullivan R, Fridman R, Ishai-Michaeli R, Sasse J et al (1987) Endothelial cell-derived basic fibroblast growth factor: synthesis and deposition into subendothelial extracellular matrix. Proc Natl Acad Sci USA 84:2292–2296

Wang Q et al. (2019) IgE-mediated mast cell activation promotes inflammation and cartilage destruction in osteoarthritis. Elife 8:e39905; doi: 10.7554/eLife.39905

Xia L, Estmuckett A, An G, Ivanciu L, McDaniel JM, Lupu F et al (2004) Defective angiogenesis and fatal embryonic haemorrhage in mice lacking core 1 - derived O-glycans. J Cell Biol 164:451–459

